# Genomic Insights into Bacterial Communities of Coenocytic Algae Using Metagenome Assembled Genomes

**DOI:** 10.64898/2026.07.19.739436

**Authors:** Gerardo Laureano, Xochitl Ramirez, Ashley Scoles, Christina Johne, Coralys M Colón, Yerlianys M. Hernández Ortiz, Jennifer Soleyman, Ramón E. Rivera Vicens, Alok Arun

## Abstract

Coenocytic algae are organisms that undergo karyokinesis without cytokinesis, resulting in multinucleated cells. Most research on bacterial communities in coenocytic algae has used 16S rRNA sequencing, primarily focusing on the order Bryopsidales of green coenocytic algae. Recent studies have analyzed metagenome-assembled genomes (MAGs) from algal hosts across multiple taxa, such as Chlorophyta, Phaeophyta, and Rhodophyta, revealing more about bacterial biosynthetic machinery and potential symbiotic relationships. Given the cosmopolitan distribution of coenocytic algae, such as *Bryopsis*, *Caulerpa*, *Codium*, and the yellow-green alga *Vaucheria*, and their unique morphology, there is a need to better understand their associated bacterial communities. To address this, filaments of *Vaucheria bursata* LB2067 were sequenced using the Illumina NovaSeq instrument, and all publicly available short– and long-read datasets from coenocytic algae were screened for MAG recovery. All recovered MAGs from both Bryopsidales and Vaucheriales showed a high dominance of Pseudomonadota at the phylum level, but no consistent patterns at the order or family levels. High completeness of specific KEGG pathways, such as bidirectional polyphosphate metabolism and riboflavin biosynthesis, was prevalent in MAGs from coenocytic orders compared to non-coenocytic ones. Notably, N-acetylglutaminylglutamine amide (NAGGN) biosynthetic gene clusters (BGCs) were found only in MAGs from coenocytic orders, whereas polysaccharide utilization loci (PULs) were present in all MAGs analyzed. These results indicate that bacterial communities associated with coenocytic algae are complex, with diverse survival strategies adapted to challenging and variable environments.

## Introduction

Eukaryotic organisms possess a specific microbiome in symbiotic relationships that can perform vital functions for the host, such as nutrient provision and protection against pathogens (Compant et al., 2019; Peixoto et al., 2021). A microbiome is a microbial community that occupies a well-defined habitat or host and performs distinct physicochemical functions (Vila Duplá 2025). The composition of a microbiome can include a wide range of microorganisms, such as bacteria, archaea, algae, and fungi. The microbiome in aquatic environments is crucial and can influence the host’s health and adaptation to environmental stress. This is significant in algae, given their symbiotic relationship with the bacterial community in the microbiota (Van Der Loos et al., 2019).

The majority of research on the composition of algal bacterial communities has focused on sequencing specific regions of the 16S rRNA gene (Singh & Reddy, 2014). This approach is widely used because the 16S rRNA gene is highly conserved, omnipresent in bacteria, and evolves slowly, making it ideal for taxonomic identification. However, if diversity is not present in the 16S rRNA region, identification fails (Lamoureux et al., 2022). A key limitation is that 16S rRNA sequencing is only for taxonomic prediction and diversity analysis (Mathieu et al., 2023). It cannot detail metabolic interactions with algae and bacteria or reveal all interactions or pathways.

In contrast to 16S rRNA sequencing, whole-genome sequencing (WGS) of the algae host enables the recovery of metagenome-assembled genomes (MAGs). This approach has proven to be a robust method for comprehending both the metabolic interactions and taxonomic identification of algal microbiomes (Meziti et al., 2021; Mirete et al., 2025). Such metabolic exchanges include algal carbon cycling, where heterotrophic bacteria, such as those in the phylum Bacteroidota, possess numerous polysaccharide-degrading enzymes and polysaccharide utilization loci (PULs) (Martens et al., 2011; Lu et al., 2023). Enzymes encoded in these gene clusters enable the microbiome to degrade host polysaccharides such as alginate, fucoidan, agar, and carrageenan (Gu et al., 2023; Lu et al., 2023; Zhang et al., 2024; Feng et al., 2022; Grondin et al., 2017). This degradation provides the bacterial community with essential building blocks for the production of metabolites, such as vitamin B12 (Balabanova et al., 2021). In contrast to PULs, Biosynthetic Gene Clusters (BGCs) are groups of genes that encode different enzymatic machinery required to produce specific metabolites (Covington et al., 2021). BGCs are critical for many organisms, particularly bacteria, as these secondary metabolites underpin essential pharmaceuticals, agrochemicals, and ecological mediators (Blin et al., 2023, 2025). Identification of BGCs and PULs enables an analysis of metabolic capabilities that cannot be determined from 16S rRNA alone.

In recent years, macroalgal bacterial communities have been recovered using metagenome-assembled genomes (MAGs) from diverse taxa, including Phaeophyceae, Rhodophyceae, and Chlorophyceae, from the genera *Nereocystis, Saccharina, Pyropia, and Ulva*, respectively. Metabolic reconstruction of these MAGs has revealed their core functional potential, particularly in carbohydrate metabolism and the degradation of host structural polysaccharides, as indicated by their high abundance (Lu et al., 2023; Wang et al., 2022; Weigel et al., 2022; Zhang et al., 2024). Additionally, these studies suggest that macroalgal-associated bacteria have the capacity to produce essential bioactive compounds and secondary metabolites, via BGCs involved in the synthesis of vitamins and phytohormones such as auxin, as well as enzymes like ACC deaminase that may help regulate host stress responses (Lu et al., 2023; Wang et al., 2022; Weigel et al., 2022).

Bacterial communities associated with yellow-green algae (Xanthophyceae), a distinct algal group characterized by xanthophyll pigments, remain poorly explored. Xanthophyceae exhibit an exceptional range of morphologies, including unicellular forms, coenocytic filaments, simple multicellular thalli, and complex multicellular thalli. Coenocytic organisms consist of multinucleate cells that lack internal cross-walls (septa), forming a continuous cytoplasm. Despite this remarkable morphological diversity, relatively little is known about the composition of their associated bacterial communities or the metabolic interactions that may differ across these growth forms. A recent study (Ochiai et al., 2025) reconstructed MAGs from the coenocytic alga *Bryopsis* sp. KO-2023 using both long– and short-read sequencing, reporting a dominance of Alpha-proteobacteria and Bacteroidota. Apart from this work, most studies of bacterial communities associated with coenocytic algae have relied primarily on 16S rRNA sequencing.

Coenocytic algae represent a unique morphological organization in which repeated nuclear divisions (karyokinesis) occur without cytokinesis, resulting in a multinucleate organism composed of a continuous cytoplasm lacking internal cross-walls. This body plan has evolved independently in several algal lineages, including the green algal genera *Caulerpa* and *Bryopsis* and the yellow-green algal genus *Vaucheria*, all of which have broad, cosmopolitan distributions. Because coenocytic algae possess a continuous cellular architecture distinct from both unicellular and multicellular forms, they provide an excellent model for investigating how algal morphology influences the composition, spatial organization, and metabolic interactions of associated bacterial communities. Analysis of bacterial communities of coenocytic algae has been dominated by 16S rRNA amplicon sequencing in *Caulerpa*, *Bryopsis,* and *Vaucheria*, revealing bacterial communities composed of Proteobacteria, Bacteroidetes, and Planctomycetes (Aires et al., 2015, 2013; Delbridge et al., 2004; Kopprio et al., 2021; Liang et al., 2019; Roth Schulze et al., 2018; Devine et al., 2012; Ochiai et al., 2025; Van Duijnhoven et al., 2024). Enzymes encoded in these gene clusters enable the microbiome to degrade host polysaccharides.

To date, the bacterial community composition of *Vaucheria* has only been reported for *V. litorea* using 16S rRNA sequencing (Devine et al., 2012). A deeper understanding of both the taxonomic composition and metabolic interactions of bacterial communities associated with *Vaucheria* species could provide valuable insights into the relationship between algal morphology and bacterial community composition. *Vaucheria* belongs to the class Xanthophyceae and is taxonomically distinct from coenocytic green algae such as *Bryopsis* and *Caulerpa* (Chlorophyta). This study aims to investigate potential associations between bacterial communities and coenocytic algal morphology through a comprehensive analysis of recovered metagenome-assembled genomes (MAGs), with a particular focus on identifying conserved metabolic functions across the surfaces of these giant single-celled algae. To this end, vegetative filaments of *Vaucheria bursata* LB2067 were sequenced using the Illumina sequencing platform. In addition, recovered MAGs were compared with those obtained from 11 publicly available whole-genome sequencing (WGS) datasets from various other coenocytic algae.

## Materials and Methods

### Algae datasets used in this study

The following genera of coenocytic algae were selected based on the availability of sequencing data in the NCBI SRA database accessed in March 2026: *Ostreobium queketti, Derbesia tenuissima, Derbesia marina, Codium adhaerens, Caulerpa lentillifera, Caulerpa ashameadii, Avrainvillea amadelpha, Penicillus capitatus, Penicillus lamourouxili, Caulerpa prolifera* and *Vaucheria bursata* (Table 1). The *Bryopsis* genus was implemented from reported MAGs recovered from *Bryopsis* KO-2023 *sp*. (Ochiai *et al.,* 2025).

**Table 1.** Summary statistics of total MAGs extraction from WGS reads of coenocytic algae by quality as high quality (HQ), good quality (GQ), and low quality (LQ).

| Organism | Order Taxonomy | Bioproject | Total MAGs | Near Complete | HQ | GQ | LQ | LR | SR | Hybrid |
| --- | --- | --- | --- | --- | --- | --- | --- | --- | --- | --- |
| *Vaucheria bursata* LB2067 | Vaucheriales | PRJNA1020787 | 11 | 0 | 10 | 1 | 0 | 0 | 11 | 0 |
| *Vaucheria bursata* | Vaucheriales | PRJNA894593 | 30 | 0 | 22 | 8 | 0 | 0 | 30 | 0 |
| *Ostreobium queketti* | Bryopsidales | PRJEB39309 | 37 | 0 | 34 | 3 | 0 | 36 | 0 | 1 |
| *Derbesia tenuissima* | Bryopsidales | PRJNA924561 | 64 | 0 | 43 | 16 | 5 | 0 | 64 | 0 |
| *Derbesia marina* | Bryopsidales | PRJNA924561 | 41 | 0 | 37 | 4 | 0 | 0 | 41 | 0 |
| *Codium adhaerens* | Bryopsidales | PRJNA924561 | 34 | 0 | 28 | 5 | 1 | 0 | 34 | 0 |
| *Caulerpa lentillifera* | Bryopsidales | PRJDB5734 | 4 | 0 | 2 | 2 | 0 | 3 | 1 | 0 |
| *Caulerpa ashmeadii* | Bryopsidales | PRJNA515488 | 6 | 0 | 6 | 0 | 0 | 0 | 6 | 0 |
| *Avrainvillea amadelpha* | Bryopsidales | PRJNA924561 | 30 | 0 | 17 | 13 | 0 | 0 | 30 | 0 |
| *Penicillus capitatus* | Bryopsidales | PRJEB87011 | 61 | 16 | 32 | 5 | 8 | 61 | 0 | 0 |
| *Penicillus lamourouxii* | Bryopsidales | PRJEB88235 | 21 | 4 | 8 | 2 | 5 | 21 | 0 | 0 |
| *Caulerpa prolifera* | Bryopsidales | PRJEB94584 | 7 | 1 | 5 | 0 | 1 | 7 | 0 | 0 |

### Culture conditions, DNA extraction, and sequencing

*V. bursata* LB2067 cultures were obtained from the UTEX Culture Collection of Algae at UT-Austin (Austin, USA). The cultures were maintained under a 16:8 light/dark photocycle at 25 °C. Isolation of genomic DNA from filaments was performed using the NucleoSpin Plant II kit (Macherey-Nagel), according to the manufacturer’s instructions. Purified genomic DNA was used for library construction following a standard Illumina library preparation protocol. Briefly, DNA was fragmented and subjected to end repair, adenylation of 3′ ends, adapter ligation, and PCR amplification to generate indexed sequencing libraries. Library concentration and insert-size distribution were evaluated prior to sequencing. Indexed libraries were pooled and sequenced on an Illumina NovaSeq instrument to produce 150-bp paired-end reads.

### Metagenome assembly and binning

Different strategies for Metagenome-Assembled Genome (MAG) recovery were performed according to the data availability for each alga: short-read (SR), long-read (LR), or hybrid (SR+LR). For datasets consisting solely of Illumina short reads (SR), the metaWRAP pipeline (v1.3) was used (Uritskiy et al., 2018), specifically the Read_QC, Assembly (with MegaHit), Binning, Bin_Refinement, and Reassembly_Bin modules. Bins with a total size exceeding 10 MB were removed to filter out potential eukaryotic contamination or assembly artifacts. All software packgages were used with default parameters unless otherwise specified. Host DNA was not removed prior to binning to support a separate, ongoing study. For datasets consisting only of long-read (LR) data (PacBio or Nanopore), the following pipeline was used: Raw long reads were assembled using Flye (v2.9.6) (Freire et al., 2022) with the ––meta option. Depending on the sequencing platform, the ––pacbio-raw, ––pacbio-hifi, or ––nano-raw flag was applied. Raw long reads were mapped to the resulting assemblies using minimap2 (v2.30) (Li 2018) with the appropriate preset (map-pb, map-hifi, or map-ont). Coverage depth across contigs was calculated using the jgi_summarize_bam_contig_depths script from MetaBAT2 (v2.18) (Kang et al., 2019). Both MaxBin2 (v2.2.7) (Wu et al., 2016) and MetaBAT2 were implemented for the binning process. The final bin set was produced by de-replicating and consolidating bins from both MaxBin2 and MetaBAT2 using DAS Tool (v1.1.7) (Sieber *et al.,* 2018) (Figure S1).

In cases where datasets were available for both LR and SR (i.e., hybrid), the processes described above were performed independently for each data type. To identify potentially identical bins between the SR and LR bin sets, FastANI (v1.34) (Jain et al., 2018) was implemented with a 95% threshold, and the resulting pairs were processed in parallel. The raw reads were mapped using minimap2 (v2.30) for LR and Bowtie2 (v2.5.4) (Langmead and Salzberg, 2012) for SR to the bins identified as identical by FastANI. The corresponding mapping files were converted to BAM format, and mapped FASTQ files were extracted using SAMtools (v1.22.1) (Li *et al.,* 2009). Reassembly was performed using Unicycler (v0.5.1) (Wick et al., 2017), and the quality of the reassembled bins was compared against the original bin pairs in terms of contamination, completeness, genome size, N50, and contig count using CheckM2 (v1.1.0) (Chklovski *et al*., 2023). The highest quality bins were selected for the final bin set (Figure S1).

### MAG quality assessment, de-replication, and taxonomic classification

The quality of the final MAGs was determined using CheckM2 (v1.1.0) to estimate genome completeness and contamination. Based on this predicted quality from CheckM2, MAGs were classified as near complete, high quality (HQ), good quality (GQ) and low quality (LQ) (Almeida et al., 2019). For HQ classification, MAGs with equal to or greater than 90% completion and equal to or less than 5% contamination were considered. MAGs classified as GQ had completion of equal or greater than 50% and equal or less than 5% contamination. In addition, LQ classification was considered by MAGs with equal or less than 50% of completion and equal or less than 10% contamination. MAGs with a completeness of at least more or equal to 90% and contamination of at most less than 5%, with only 1 contig, were classified as near complete. MAGs with a contamination level of 10% or higher were removed from the subsequent analysis, regardless of completeness; only 1 MAG was removed. De-replication of MAGs was performed to remove replicated genomes from the dataset by implementing Galah (v0.4.2) (https://github.com/wwood/galah) for clustering. The CheckM2 output was then used to manually select the best representative genome from each cluster based on contamination, completeness, genome size, N50, and contig count.

Taxonomic classifications were conducted primarily using GTDB-Tk (2.6.1+galaxy0) (Genome Taxonomy Database Toolkit) (Chaumeil et al., 2022) using the database version 224. Using classifications derived from GTDB-Tk, taxonomic lineages were assigned. Barrnap (v0.9) (https://github.com/tseemann/barrnap) was used to identify all rRNA sequences present in MAGs. QIIME 2 (v2025.10) (Bolyen et al., 2019) was used with the SILVA 138.2 database (Quast et al., 2012) for taxonomic prediction of the extracted 16S sequences for validation of genus and species of GTDB-tk results. For this, only SILVA similarity scores of equal to or greater than 94.5% and 98.7% were used for genus and species, respectively.

### Structural analysis, functional analysis and BGCs discovery

Structural annotation was performed using Bakta (v1.9.11) (Schwengers et al., 2021) with the –– metagenome option. For comprehensive functional annotation, genes were processed using eggNOG-mapper (v2.1.13) (Cantalapiedra et al., 2021). KEGGaNOG (v1.1.19) (https://github.com/iliapopov17/KEGGaNOG) was implemented to evaluate the completeness of KEGG pathways present in each MAG using the output from eggNOG-mapper. AntiSMASH (v8.0.4) (Blin et al., 2025) was used to discover Biosynthetic Gene Clusters (BGCs), and BiG-SCAPE (v2.0.0) (https://github.com/medema-group/BiG-SCAPE) was used to classify and count BGC families present throughout the algal community (Figure S2).

### PULs identification and classification

PULs were broadly defined as genomic regions containing at least three genes coding for CAZymes, sulfatases, or transport proteins, following the criteria reported analysis in the microbiome analysis of co-located green, brown, and red algae (Lu et al., 2023). These loci were assigned to four compositional categories: PULs comprising CAZymes and a *susCD* pair; CAZyme-rich gene clusters (CGCs) lacking transporter genes; PUL-like clusters containing CAZymes and a TonB-dependent receptor; and *susCD* loci without detectable CAZymes. Genes containing SusC/D-like proteins were predicted with HMMER with an E-value cutoff of 1 × 10^-10^ against Pfam and TIGRFAM profiles. PFAM profiles used to identify sulfatase were PF00884, for SusD-like, PF07715, PF07980, PF12741, PF14322, PF12771, and TonB-dependent receptor were PF00593. The TIGRFAM profiles were TIGR04056 TIGR01352, TIGR01778, TIGR01779, TIGR01782, TIGR01783, TIGR01785, TIGR01786, TIGR02796, TIGR02797, TIGR02803, TIGR02804, TIGR02805, TIGR04057 (Table 2). CAZymes were identified with run_dbCAN (https://github.com/bcb-unl/run_dbcan) (v5.2.1) with the dbCAN3 database (Zheng et al., 2023).

**Table 2.** Summary of Pfam and TIGRFAM profiles utilized in PULs identification based on locus category.

| Locus Category | Defining Features | Pfam Profiles | TIGRFAM Profiles |
| --- | --- | --- | --- |
| PULs | CAZymes + susCD pair | SusD-like: PF07715, PF07980, PF12741, PF14322, PF12771 | SusC/D-like: TIGR04056, TIGR01352, TIGR01778, TIGR01779, TIGR01782, TIGR01783, TIGR01785, TIGR01786, |
|  |  |  | TIGR02796, TIGR02797, TIGR02803,<br>TIGR02804, TIGR02805, TIGR04057 |
| PUL-like clusters | CAZymes + TonB-dependent receptor | TonB-dependent receptor: PF00593 | N/A |
| CGCs | CAZymes | N/A | N/A |

### Comparison with Non-Coenocytic Algae

MAGs from *Ulva* sp. and *Saccharina* sp. of the orders Ulavales and Laminariales were used for comparative analysis of BGCs, PULs and KEGG pathways analysis, with the MAGs recovered from coenocytic algae. These MAGs were obtained from reported microbiome analysis of co-located green, brown, and red algae (Lu et al., 2023). The MAGs from non-coenocytic algae were subjected to the same methodology as the MAGs from coenocytic algae in BGCs, PULs, and KEGG pathways. Statistical comparisons of pathway completeness between coenocytic and non-coenocytic MAGs were conducted in R (v4.5.0). Only pathways detected in at least 50% of MAGs in both groups were included in the analysis. For each pathway, a one-tailed Mann– Whitney U test (alternative hypothesis: coenocytic > control) was performed, and p-values were adjusted for multiple testing using the Benjamini–Hochberg method (adjusted *p* < 0.05). Effect sizes were calculated as rank-biserial correlations (*r*) and classified as negligible (|*r*| < 0.1), small (0.1 ≤ |*r*| < 0.3), medium (0.3 ≤ |*r*| < 0.5), or large (|*r*| ≥ 0.5). Data processing and visualization were performed using the R packages dplyr (v1.2.0), tidyr (v1.3.2), ggplot2 (v4.0.2), and rstatix (v0.7.3).

## Results

### Coenocytic algae MAG quality and size distribution

The MAGs recovered from the coenocytic WGS reads, and *V. bursata* LB2067 showed a total of 346 MAGs (Table 1, Table S1 & Figure 1H). A total of 262 MAGs were recovered from 11 algae of the order Bryopsidales; 16 were near complete, 179 were HQ, 61 were GQ, and 6 were LQ. From the 2 algae of the order Vaucheriales, a total of 41 MAGs were recovered, 22 were HQ, and 8 were GQ (Table 1 & Table S2). A total of 132 MAGs were from SR, 127 from LR, and 1 hybrid. MAGs from order Bryopsidales and Vaucheriales showed similar quality in terms of N50, fragmentation, and genome size, with MAGs from Bryopsidales having a higher value in the metrics, respectively (Figure 1D-F). The recovery of a high number of HQ MAGs confirms that the proposed methodology is a robust approach for obtaining high-quality genomes from the WGS of both *Bryopsidales* and *Vaucheriales.* This is further illustrated in the completion and contamination scatter plot (Figure 1G). In contrast to the quality of MAGs recovered, the genome size distribution between both orders shows high variations, where MAGs from Bryopsidales algae have a genome size distribution between 1 and 10 Mbps with reduced genomic elements of less than 1 Mbp (Table S2). MAGs from Vaucheriales have a lower distribution of 3 to 9 Mbps with a mean genome size of approximately 5 Mbps (Figure 1F).

**Figure 1.**
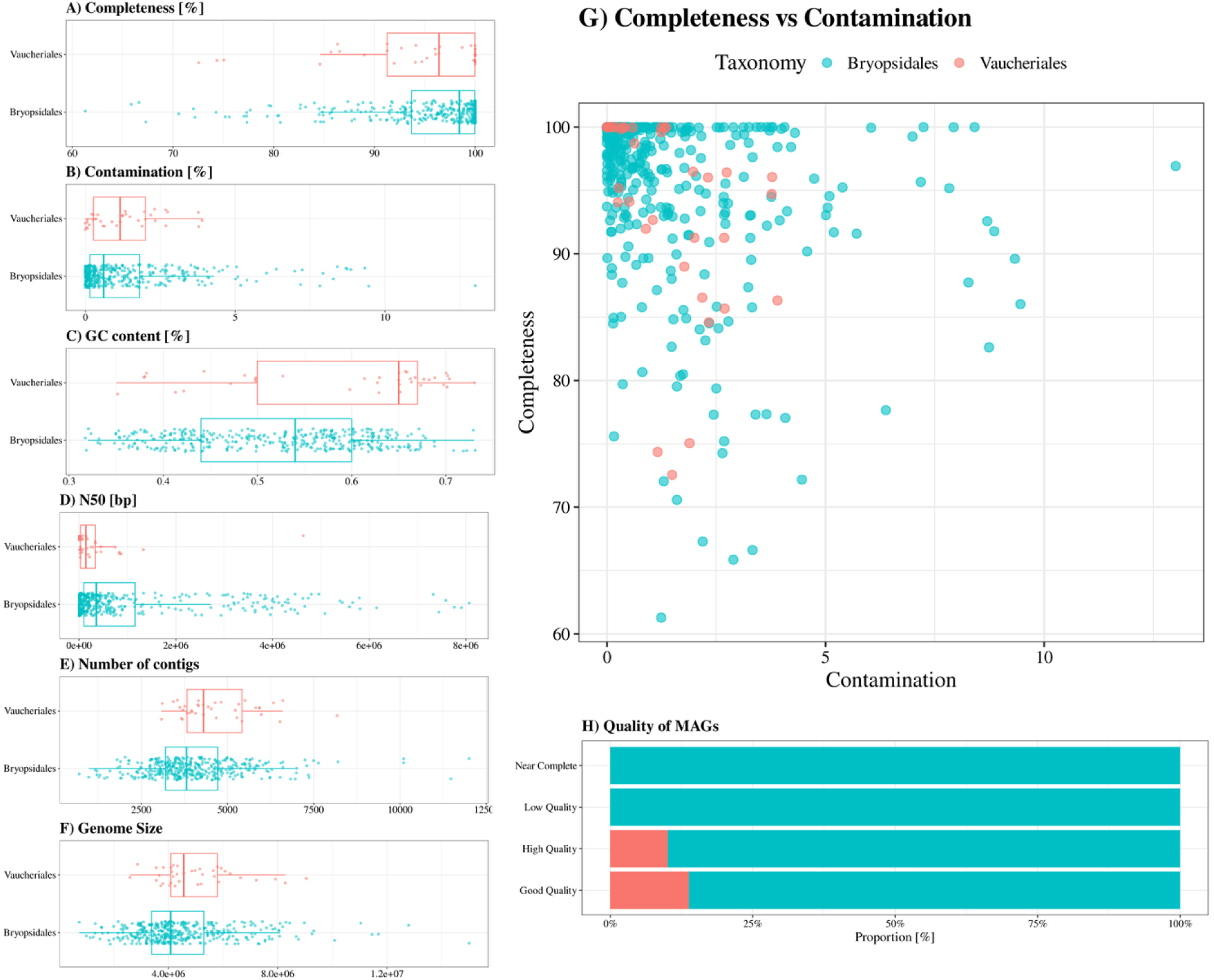
Quality metrics and genome size distributions of MAGs recovered from coenocytic algae metagenomes. (A) Distribution of genome completeness estimated by CheckM2 across coenocytic alga orders. (B) Distribution of contamination levels. (C) GC content distribution. (D) Assembly N50 distribution. (E) Number of contigs per MAG. (F) Total genome size distribution. (G) Scatter plot illustrating the relationship between completeness and contamination. (H) Categorization of MAGs based on quality tiers.

### Taxonomic Composition of the Coenocytic Algae

The taxonomic profiles of the recovered MAGs from coenocytic orders Bryopsidales and Vaucheriales, displayed broad similarities at the phylum level but diverge significantly at different taxonomic resolutions (Figures 2A & 2B). At the phylum level, both lineages are heavily dominated by Pseudomonadota and Bacteroidota. However, the extent of this dominance varies; in the MAGs from Vaucheriales, these two phyla collectively comprise 84.4% of the microbiome, whereas in the MAGs from Bryopsidales, they account for 72.9%, with the latter showing a notable enrichment of Cyanobacteriota and Planctomycetota (Table S3 & S4).

**Figure 2.**
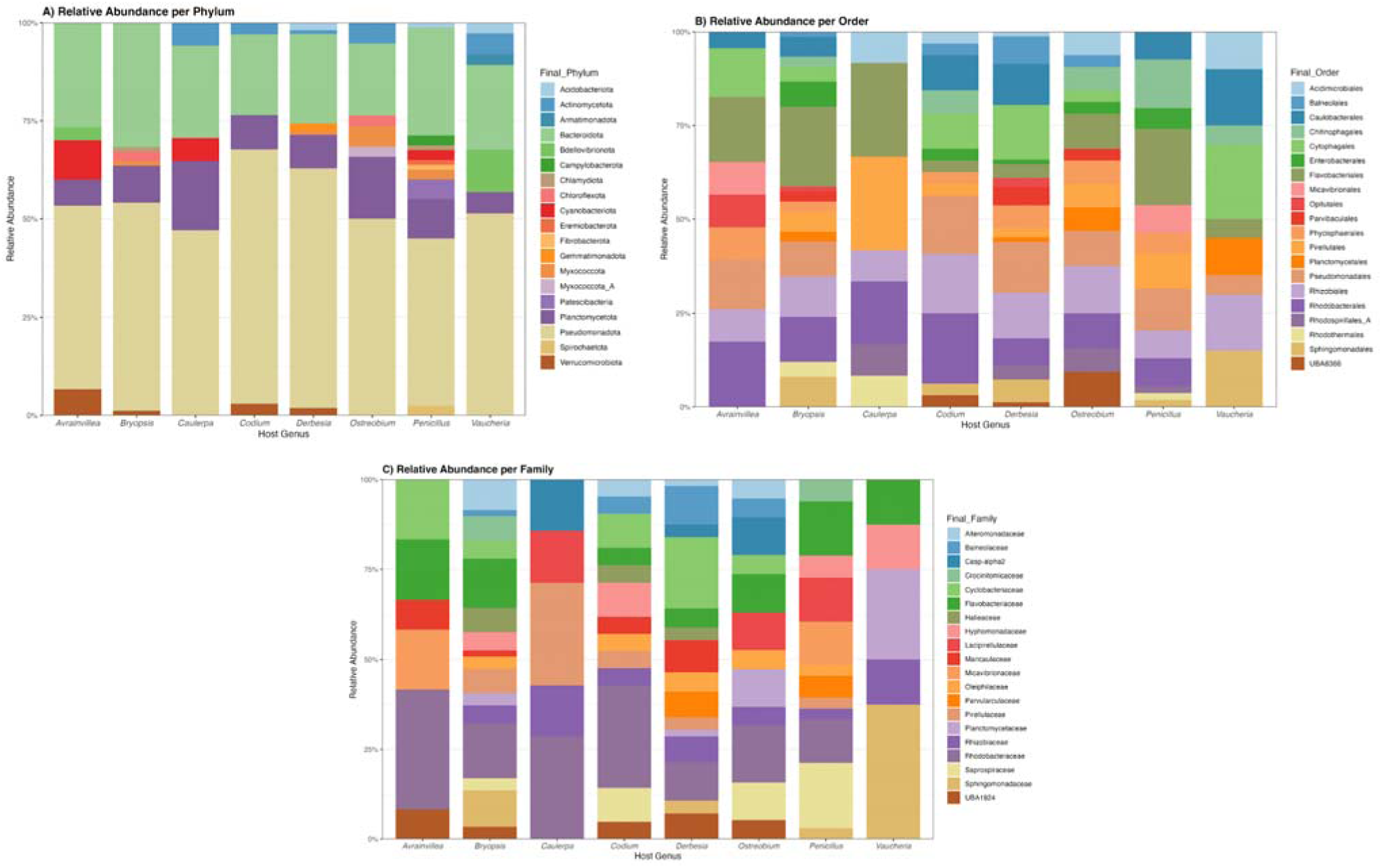
Taxonomical comparison between algae from the coenocytic order Bryopsidales and Vaucheriales by GTDB-tk prediction and SILVA 138 99% OTUs full-length sequences database. (A) Distribution of the top phylum through MAGs recovered from coenocytic algae. (B) Distribution of top order. **(C) Distribution of top family-level.**

At the order level (Figure 2B), this divergence becomes more pronounced. The MAGs from the Vaucheriales are characterized by an even distribution with Sphingomonadales, Burkholderiales, and Rhizobiales each contributing 11.1% (cumulatively 33.3%). In contrast, the Bryopsidales MAGs composition is co-dominated by Rhizobiales and Flavobacteriales (both 11.5%), followed by Rhodobacterales (10.7%) and Pirellulales (7.6%). The Flavobacteriaceae is also present in the Vaucheriales MAGs, but in lower concentration. The high biological diversity in taxonomy prediction is at family level (Figure 2C). The MAGs from Vaucheriales retain a consistent taxonomy distribution of Sphingomonadaceae 11.1%, Burkholderiaceae 8.9%, and Rhizobiaceae 8.9%, with the notable addition of Bacteriovoracaceae 6.7%. In contrast, the Bryopsidales display high specificity for families associated with complex host interactions, including Rhodobacteraceae, Flavobacteriaceae, Pirellulaceae, and the Oleiphilaceae. Furthermore, GTDB-Tk classification revealed a significant accumulation of novel taxonomic diversity in the Bryopsidales dataset, with more than 50% of their MAGs classified as putative novel genera.

### KEGG pathways from recovered MAGs of coenocytic orders

KEGG pathways across the MAGs from the coenocytic orders Bryopsidales and Vaucheriales, when compared with MAGs from non-coenocytic algae (*Ulva*, *Saccharina*), revealed a conserved non-photosynthetic bacteria core, with all lineages encoding consistent pathways for amino acid biosynthesis, glycolysis, and the TCA cycle. Depending on the algal order from which the MAGs were recovered, distinct KEGG pathways were observed with varying levels of completeness. The MAGs recovered from the Bryopsidales show specific pathway presence, such as nitrogen fixation, in MAGs from the *Derbesia* and *Bryopsis* KO-2023 sp. Conversely, MAGs in *Penicillus* and *Avrainvillea* are partially complete, given the value of KEGGaNOG results (Table S5) in the same pathways. In contrast, MAGs from the Vaucheriales exhibited complete nitrogenase modules co-occurring with Cytochrome *bd* ubiquinol oxidase and Polyhydroxybutyrate (PHB) synthesis. Furthermore, motility profiles differed sharply; MAGs from non-coenocytic associates possessed complete flagellar assembly pathways, whereas the MAGs from Bryopsidales uniquely retained a specific, partial flagellar/type III secretion system signature indicative of a stable sessile lifestyle.

A principal component analysis (PCA) of bacterial KEGG pathways revealed a broad functional similarity across the majority of MAGs in both, non-coenocytic and coenocytic algae (Figure S4). However, the MAGs from Vaucheriales displayed a distinct separation from the Bryopsidales. This divergence was most pronounced in the LB2067 dataset, which formed a cluster completely distinct from the main group, in contrast to MAGs from the other *V. bursata* datasets, which showed only moderate divergence and remained closer to the central functional array. Within the Bryopsidales, while most samples clustered tightly, specific *Caulerpa* species displayed notable variance, diverging from the major group arrays (Figure S4).

A Mann-Whitney U test, non-parametric and adjusted for multiple testing using the Benjamini-Hochberg method, was used to identify pathways with significantly different completeness between coenocytic and non-coenocytic MAGs (padj < 0.05). Of the 56 pathways shared between the two groups, 46 were statistically significant (Table S8). Among these, 18 had higher completeness of coenocytic MAGs, including pathways such as protein secretion (Sec-SRP), motility (Flagellum, Chemotaxis), and central carbon metabolism (Glycolysis, TCA Cycle). The remaining 28 pathways were more complete in non-coenocytic controls, such as amino acid biosynthesis (Leucine, Glutamine, Arginine), energy metabolism (F-type ATPase, Cytochrome c oxidase), and cofactor biosynthesis (Riboflavin biosynthesis).

### BGCs distribution from recovered MAGs of coenocytic order

A comparison of the distribution of identified biosynthetic gene clusters (BGCs) shows that MAGs recovered from the Bryopsidales contained a substantially higher number of BGCs than those from the Vaucheriales, Ulvales, and Laminariales (Figure 5A). Notably, MAGs associated with Ulvales and Vaucheriales exhibited a high BGC count relative to the small number of algal hosts from which they were recovered, one and two, respectively (Figure 5A). Regarding BGC types, the majority were conserved between coenocytic and non-coenocytic algae, particularly terpenes, terpene precursors, and T1PKS (Figure 5B & Table S6). The N-acetylglutaminylglutamine amide (NAGGN) BGC type was detected only in MAGs from coenocytic algae and was absent from non-coenocytic hosts. Its distribution among the coenocytic taxa was not uniform, as NAGGN was not detected in *Avrainvillea*, *Caulerpa*, and *Codium* species. The MAGs with NAGGN were assigned almost entirely to the Pseudomonadota and were distributed across several bacterial orders rather than a single lineage.

The MAGs from the non-coenocytic algae, specifically *Ulva* sp., uniquely possessed T2PKS and PUFA clusters. In contrast, the MAGs from coenocytic algae were characterized by the exclusive presence of Butyrolactone and NAGGN clusters. Butyrolactone was specific to the MAGs from Bryopsidales, species *Bryopsis* KO-2023 (Table S5). The NAGGN BGC’s were more widely distributed among MAGs from coenocytic algae, with the highest prevalence in *Ostreobium* (33%) and *Derbesia* (32%), followed by *Bryopsis* (16%), and *Vaucheria* and *Penicillus* (8%). Despite the evolutionary distance between Vaucheriales and Bryopsidales, both shared the usage of NAGGN clusters. However, the MAGs from Vaucheriales were distinct, exclusively harboring clusters such as phosphonate-like, NRP-metallophore-NRPS-T1PKS, and various complex hybrid PKS/NRPS combinations.

### PULs distribution

The polysaccharide utilization potential of the bacterial community was assessed using MAGs to identify Polysaccharide Utilization Loci (PULs), PUL-like, and CAZyme Gene Clusters (CGCs). Through MAGs from coenocytic and non-coenocytic distinct distribution patterns were observed across all MAGs (Figure 6 & Table S7). The analysis revealed a marked predominance of PULs and PUL-like compared to CGCs across the MAGs from algae of both coenocytic and non-coenocytic orders. As shown in Figure 6, the frequency of identified loci was consistently higher for PULs, particularly within the Bryopsidales MAGs, which exhibited the highest absolute counts of PULs and PUL-like. While CGCs were present across all groups, their overall abundance was notably lower than that of the PUL types.

## Discussion

### Taxonomy classification and quality

The majority of MAGs recovered from the WGS reads of the coenocytic algae were classified from high to good quality with a low level of contamination and high completeness (Figure 1 G & Table S2). This high yield of MAGs demonstrates the viability of recovering bacterial community from host tissues, aligning with recent successes in both targeted coenocytic algal studies (Ochiai et al., 2025) and broad scale macroalgal metagenomic surveys (Lu et al., 2023). The predominance of MAGs recovered from SR was classified as good quality and high quality, in contrast to LR, which were classified predominantly as high quality and near complete (Figure S3). Both sequencing methods yielded LQ MAGs; the LR assemblies showed less fragmentation, with fewer contigs and higher N50 values than the SR assemblies (Figure S3). This result is consistent with Orellana et al., (2023), who reported that short-read sequencing often recovers a higher total number of genomes due to greater sequencing depth, but long-read approaches yield a significantly higher proportion of high-quality MAGs. They reported that LR MAGs were composed of approximately 10 times fewer contigs and possessed N50 values more than 11 times higher than SR MAGs. Our results show that the workflow utilized for MAGs recovery was able to recover HQ MAGs from both SR and LR of WGS.

The distribution of taxa in both coenocytic orders, Bryopsidales and Vaucheriales, showed a direct similarity at the phylum level, characterized by a consistently high percentage of Pseudomonadota and Bacteroidota. In Bryopsidales, these two phyla constitute 72.9% of the recovered MAGs (44.7% Pseudomonadota and 28.2% Bacteroidota), while in the Vaucheriales, it accounts for 84.4% of the community (64.4% Pseudomonadota and 20.0% Bacteroidota) (Figure 2A). These findings align with previous research on Bryopsidales in which the community of *Caulerpa lentillifera* was dominated by Pseudomonadota (syn. Proteobacteria) at 52.1% and Bacteroidota (syn. Bacteroidetes) at 13.5% (Liang et al., 2019). The dominance of Pseudomonadota observed in the *V. bursata* MAGs aligns with previous 16S rRNA sequencing of *V. litorea*, which reported an 88.4% relative abundance of this phylum. However, *V. bursata* exhibited a notably higher proportion of Bacteroidota (20.0%) compared to the 7.3% observed in *V. litorea* (Devine et al., 2012). This could indicate a difference in species recruitment, though this variance may also be influenced by the differences between WGS binning and 16S rRNA sequencing. The co-dominance of Pseudomonadota and Bacteroidota is not unique to coenocytic algae but has been reported across diverse algal lineages, such as in the red alga *Pyropia haitanensis*, comprising approximately 65% of the sequenced microbiome (Wang et al., 2022). Furthermore, a major comparative across *Ulva* sp., *Grateloupia sp*., *Gelidium sp*., and *Saccharina sp*. found that core genera belonging to Pseudomonadota and Bacteroidota dominated the algal surface, comprising 51.1% of the total bacterial community compared to only 5.7% in the surrounding seawater and 1.5% in the sediment (Lu et al., 2023). The phylum distribution of MAGs between coenocytic algae is consistent with other non-coenocytic marine algae with dominance of Pseudomonadota and Bacteroidota, even in the case of *V. bursata* LB2067, which resides in freshwater.

In contrast to the phylum level, at the order level, MAGs from Bryopsidales show a specific dominance by Rhizobiales, Rhodobacterales, and Flavobacteriales across all algae, and an almost dominant position of Pseudomonadales (Figure 2B) was observed. Conversely, MAGs from Vaucheriales show a distribution of Sphingomonadales, Burkholderiales, and Rhizobiales, each contributing 11.1% (Figure 2B). In Bryopsidales, this dominance of Rhodobacterales and Rhizobiales has also been reported in *Caulerpa lentillifera,* accounting for approximately 22.0% of the bacterial community (Liang et al., 2019). Taxonomic profiling of the *V. bursata* MAGs revealed high abundances of Rhizobiales and Burkholderiales (Figure 2B & Table S4), alongside a lower abundance of Sphingomonadales (Figure 2B), compared to the known composition of *V. litorea* (Devine et al., 2012). Furthermore, while *V. litorea* harbors a high abundance of Rhodobacterales, this order was absent from the *V. bursata* MAGs. These distinctions could indicate a divergence between the bacterial communities associated with these two *Vaucheria* species.

At the family level, there is more diversity among all MAGs from coenocytic algae for both Bryopsidales and Vaucheriales (Figure 2C). Although in Bryopsidales, Flavobacteriales are present in *Bryopsis* KO 2023 sp. species and the *Penicillus* genus. In contrast, the family Flavobacteriaceae from the order Flavobacteriales is present across both Bryopsidales and Vaucheriales. In 2011, a bacterial endophyte from the Flavobacteriales order was reported to be present in *Bryopsis*. Given the major presence of the Flavobacteriaceae family across both Bryopsidales and Vaucheriales (Table S4), could indicate that a member of this order could be a potential endophyte or symbiont associated with coenocytic algae (Hollants et al., 2011). Both MAGs from Bryopsidales and Vaucheriales show a high proportion of potentially novel taxa of more than 50% and 25%, respectively, according to GTDB-tk prediction with SILVA integration for validation.

Together, all these findings highlight complex biological interactions; notably, the overall taxonomic distribution observed in these MAGs remains inextricably linked to the abiotic environmental conditions, such as salinity and pH, surrounding the algae prior to sequencing. Consequently, the taxonomic profiles generated here require further validation using complementary methodologies.

### Metabolic KEGG pathways distribution

An evaluation of KEGG metabolic pathways across the MAGs recovered from coenocytic and non-coenocytic algae reveals that core bacterial metabolism remains highly conserved (Figure 3). Specifically, pathways for glycine, cysteine, and lysine biosynthesis, as well as glycolysis and the TCA Cycle, show high completeness across all MAGs (Figure 4). Among the pathways significantly enriched in coenocytic-associated MAGs, the Sec-SRP protein secretion system showed the strongest signal (Table S8), suggesting that bacteria associated with coenocytic algae may possess enhanced secretory capacity, potentially reflecting a more intimate interaction with the host environment. Conversely, bidirectional polyphosphate (polyP) metabolism and riboflavin biosynthesis were significantly more complete in non-coenocytic controls (Mann-Whitney U, padj < 0.05), although both pathways remained highly prevalent across coenocytic MAGs, with completeness values reaching 90–100% in Vaucheriales-associated MAGs. The high prevalence of the polyP pathway is notable, as it is often described as an indicator of environmental durability typically associated with a free-living lifestyle (Wang et al., 2018), suggesting that coenocytic-associated microbiomes retain broad survival capacity despite their host association. While overall pathway completeness patterns suggested broad functional differences between coenocytic and non-coenocytic-associated bacteria, a nonparametric Mann-Whitney U test was performed to identify specifically enriched pathways in the coenocytic-associated MAGs. This analysis revealed 18 metabolic functions with significantly higher completeness. The enrichment of traits such as protein secretion (Sec-SRP) and motility (flagella, chemotaxis) may reflect ecological adaptations of these bacteria to the coenocytic host environments (Table S8). Conversely, the higher completeness observed in foundational pathways, such as central carbon metabolism (glycolysis, TCA cycle), is likely driven by methodological biases. Specifically, the control MAGs were derived exclusively from short-read assemblies, whereas 41% of the coenocytic MAGs benefited from long-read sequencing. The higher genomic contiguity inherent in long-read data likely inflated these estimates of foundational pathway completeness.

**Figure 3.**
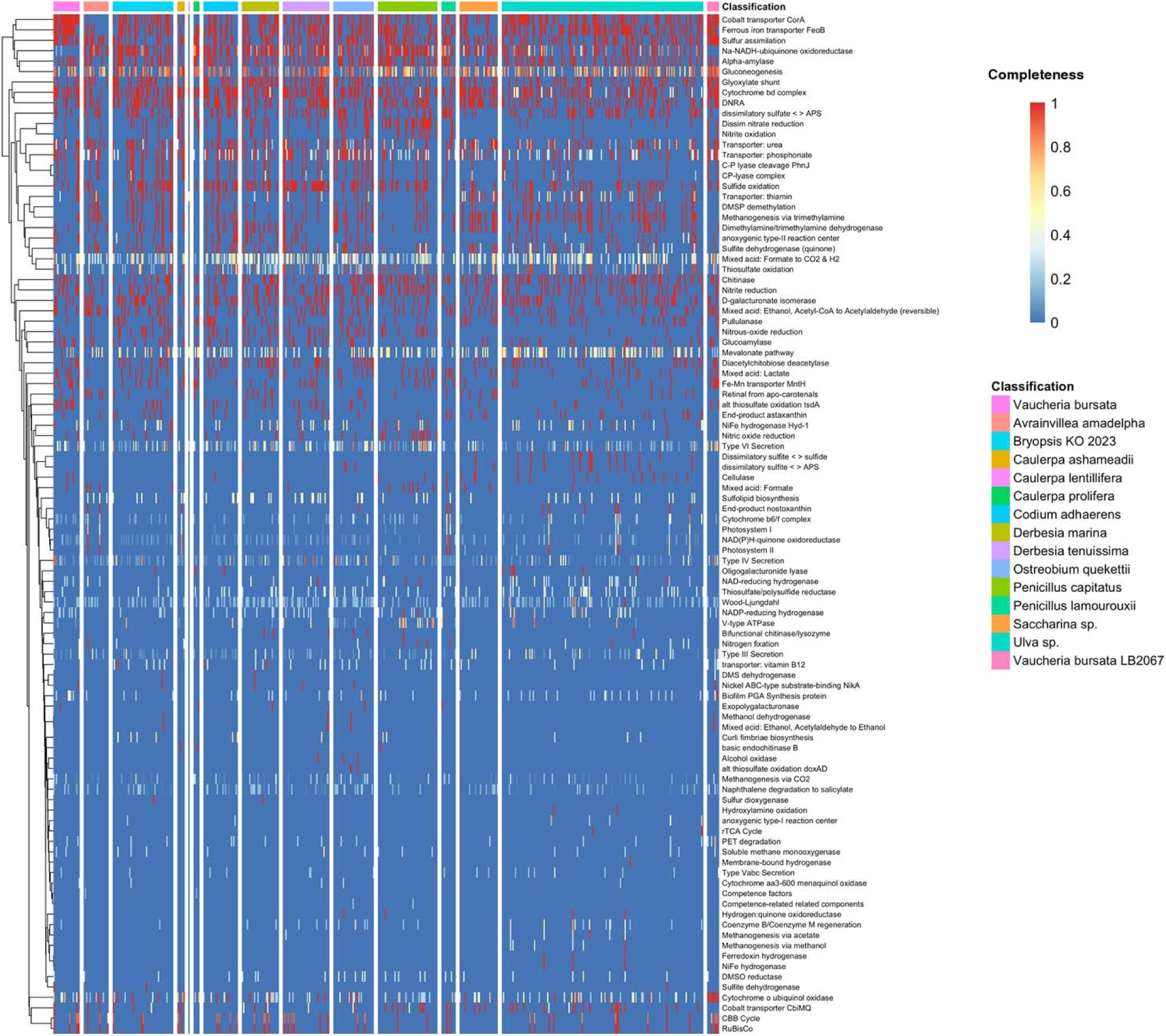
Metabolic functional profiling of bacterial communities associated with coenocytic and non-coenocytic algae. This heatmap, generated using NOG, illustrates the non-shared metabolic pathways across bacterial communities associated with coenocytic and non-coenocytic algae. The color gradient represents pathway completeness, ranging from 0 (absent, blue) to 1 (complete, red). Samples are grouped by host algal species, as indicated by the classification at the top. Hierarchical clustering highlights similarities in functional profiles among both samples and metabolic pathways.

**Figure 4.**
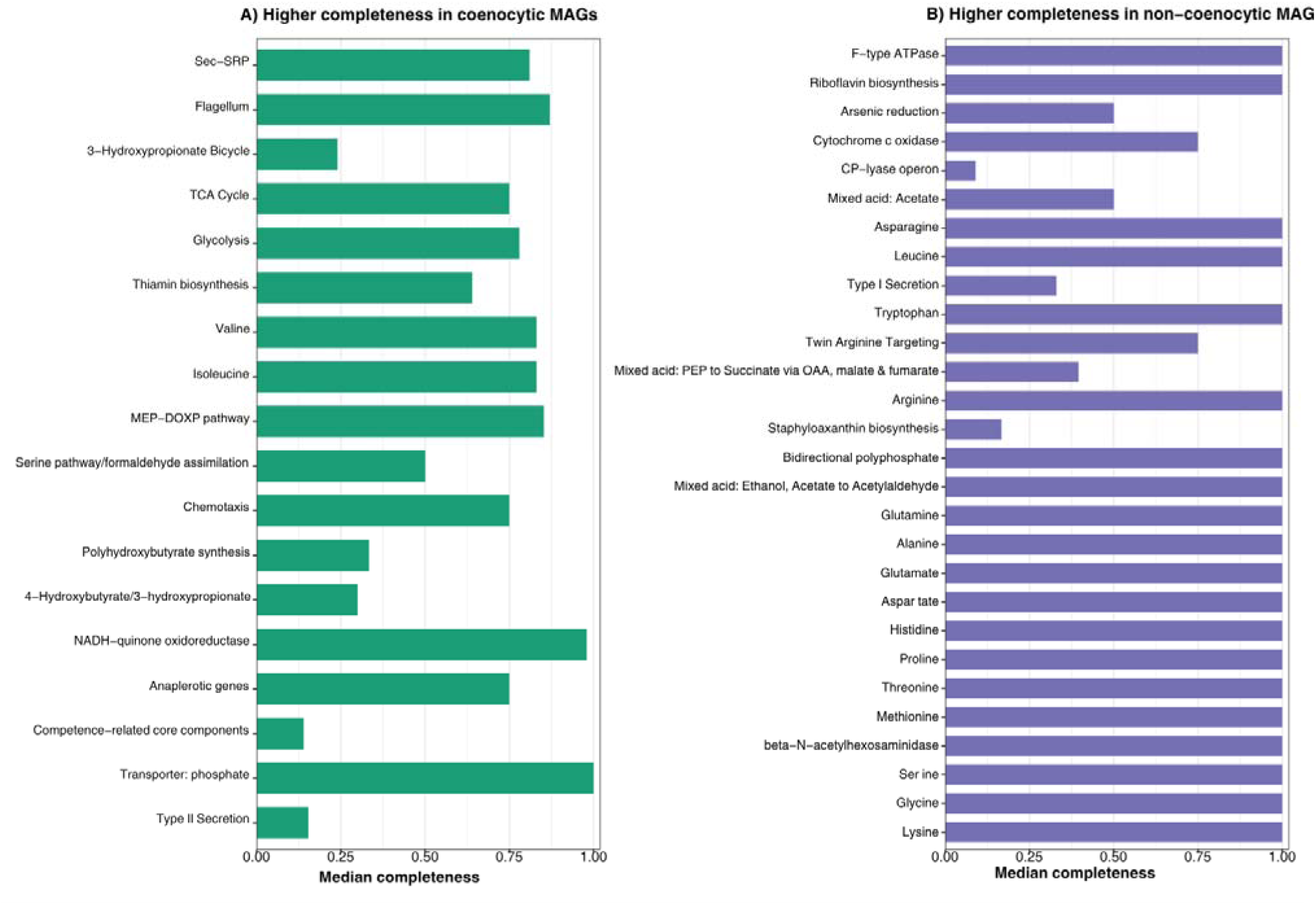
Metabolic pathways with significantly different completeness between coenocytic and non-coenocytic associated MAGs (Mann-Whitney U, padj < 0.05), ordered by effect size. (A) Pathways with significantly higher completeness in coenocytic MAGs. (B) Pathways with significantly higher completeness in non-coenocytic MAGs.

The Polyhydroxybutyrate (PHB) synthesis was significantly enriched in coenocytic-associated MAGs compared to non-coenocytic controls (Figure 4B). Within the coenocytic group, all MAGs from Vaucheriales showed a consistent completeness of 83% for the PHB synthesis pathway. Conversely, 46.9% of MAGs from non-coenocytic algae showed low completeness in this pathway, with values of 15.4% and 6.2% for Ulvales and Laminarales, respectively (Table S5). Various studies have reported that PHB synthesis serves as a critical survival mechanism, providing a carbon reserve through the accumulation of PHB polymer (Timmis 2010; Verlinden et al., 2007; Müller-Santos et al., 2021). The characteristics of PHB synthesis have been demonstrated in several bacterial species, such as *Agromyces indicus*. In 2021, PHB synthesis was optimized for *Agromyces indicus* using response surface methodology (RSM). Researchers concluded that the subsequent decrease in PHB was in direct response to a survival mechanism for the insufficient source of carbon and nitrogen in one of the conditions of the RSM analysis (Adnan et al., 2022). Given the higher prevalence of MAGs with pathway completeness for PHB synthesis (Figure 4A), suggest that the Vaucheriales bacterial community exhibits mechanisms for growth in environments with a high density of organisms, potentially as algae mats. Throughout all MAGs from the coenocytic and non-coenocytic algae, notably only one MAG from *V. bursata* LB2067 showed a complete pathway with Biofilm PGA. The MAG was taxonomically predicted by GTDB-tk to be *Achromobacter aegrifaciens*. Notably, it is not present in the MAGs from the other reported *V. bursata* algae, which, in contrast to *V. bursata* LB2067, is not a laboratory culture. This suggests that the *A. aegrifaciens* MAG is a potential lab-induced acquisition.

### BGC’s & PULs distribution

The BGCs identified in MAGs across the coenocytic and non-coenocytic orders (Bryopsidales, Vaucheriales, Ulvales, and Laminariales) were dominated by terpenes, terpene precursors, and T1PKS (Figure 5B). MAGs from Bryopsidales exhibited the highest number of BGCs, exceeding 1,500 and substantially more than any other order (Figure 5A), followed by Ulvales (∼500) and Vaucheriales (>200). For Vaucheriales MAGs, this abundance highlights a high density of BGC types within the bacterial community, given that the dataset contains only 29 non-redundant MAGs. MAGs from the analyzed algae of the Bryopsidales and Vaucheriales contain the NAGGN BGC type, in contrast to MAGs from their non-coenocytic counterparts (Figure 5B). Its distribution among the coenocytic taxa was not detected in *Avrainvillea*, *Caulerpa*, or *Codium* species. The MAGs possessing NAGGN BGC were assigned to the Pseudomonadota, spanning several orders including the Rhizobiales, Pseudomonadales, Caulobacterales, Nevskiales, Rhodospirillales, and Acidimicrobiales. The NAGGN BGCs are associated with osmoprotection mechanisms via the integration of compatible solutes, enabling bacteria to regulate internal osmotic pressure (Sagot et al., 2010). Together with the KEGG pathway analysis of the MAGs, this could indicate that bacterial communities in coenocytic algae are capable of surviving in complex environments, from permanently salty oceans to soil and the interior of plant or animal hosts (Sagot et al., 2010)). Regarding carbohydrate-active gene systems, MAGs from coenocytic algae showed a higher abundance of PULs and PUL-like systems than CGCs (Figure 6). While canonical PULs are characteristic of Bacteroidota, PUL-like clusters are more broadly distributed across bacterial lineages, including Pseudomonadota (Martens et al., 2011). The predominance of PUL-like systems could therefore be consistent with the high relative abundance of Pseudomonadota in these communities (Figure 2A). This may suggest a community-wide reliance on complex-carbohydrate-degradation machinery in the bacterial community of coenocytic algae. Notably, MAGs tend to be more fragmented. Because PULs and PUL-like systems are defined over co-located arrays of genes, assembly fragmentation can split these loci across contigs, so that they may be recovered as shorter clusters than are present. Similar limitations of MAG-based gene-cluster prediction have been noted previously (Lu et al., 2023).

**Figure 5.**
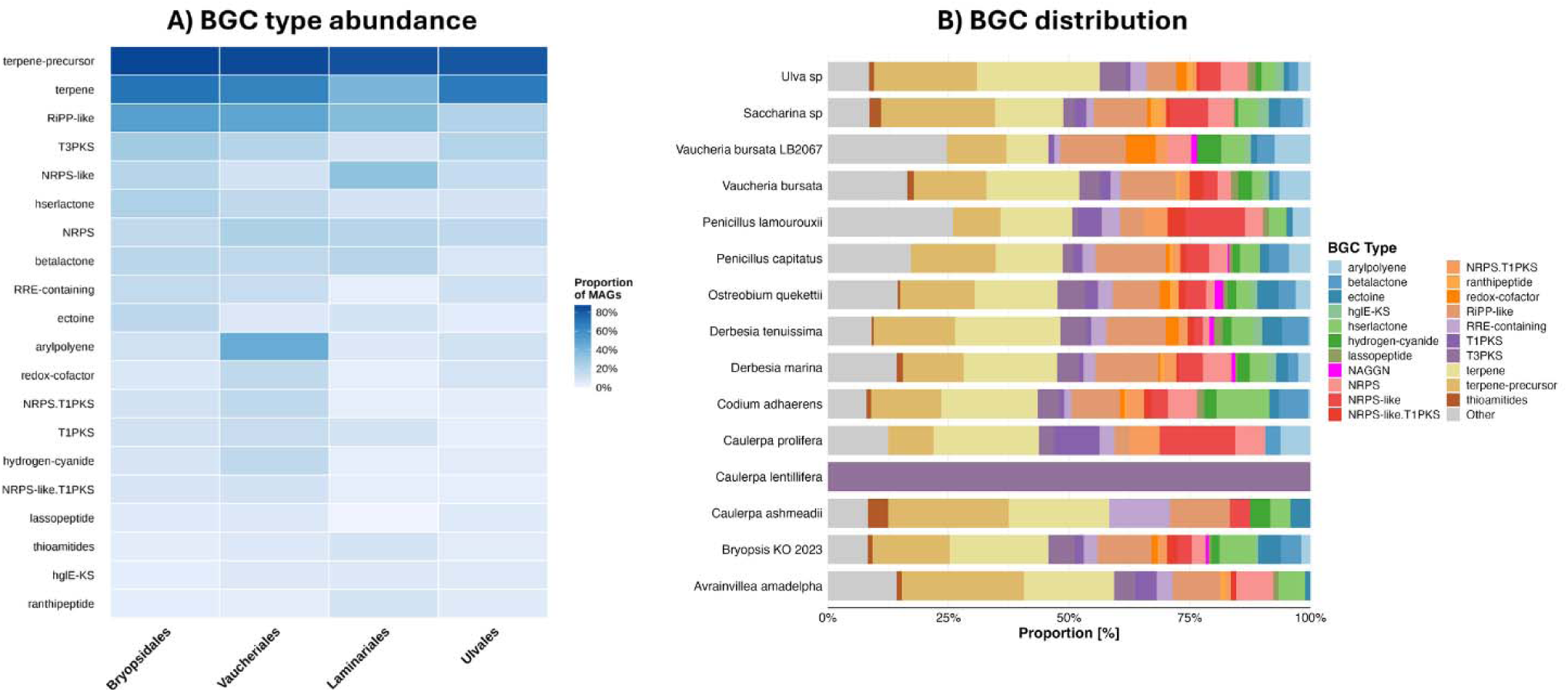
BGCs distribution through MAGs of coenocytic and non-coenocytic orders identified by antismash v8.0.4. (A) BGCs type abundance between MAGs per Order. (B) frequency distribution of the 20 most abundant BGC types across the MAGs.

**Figure 6.**
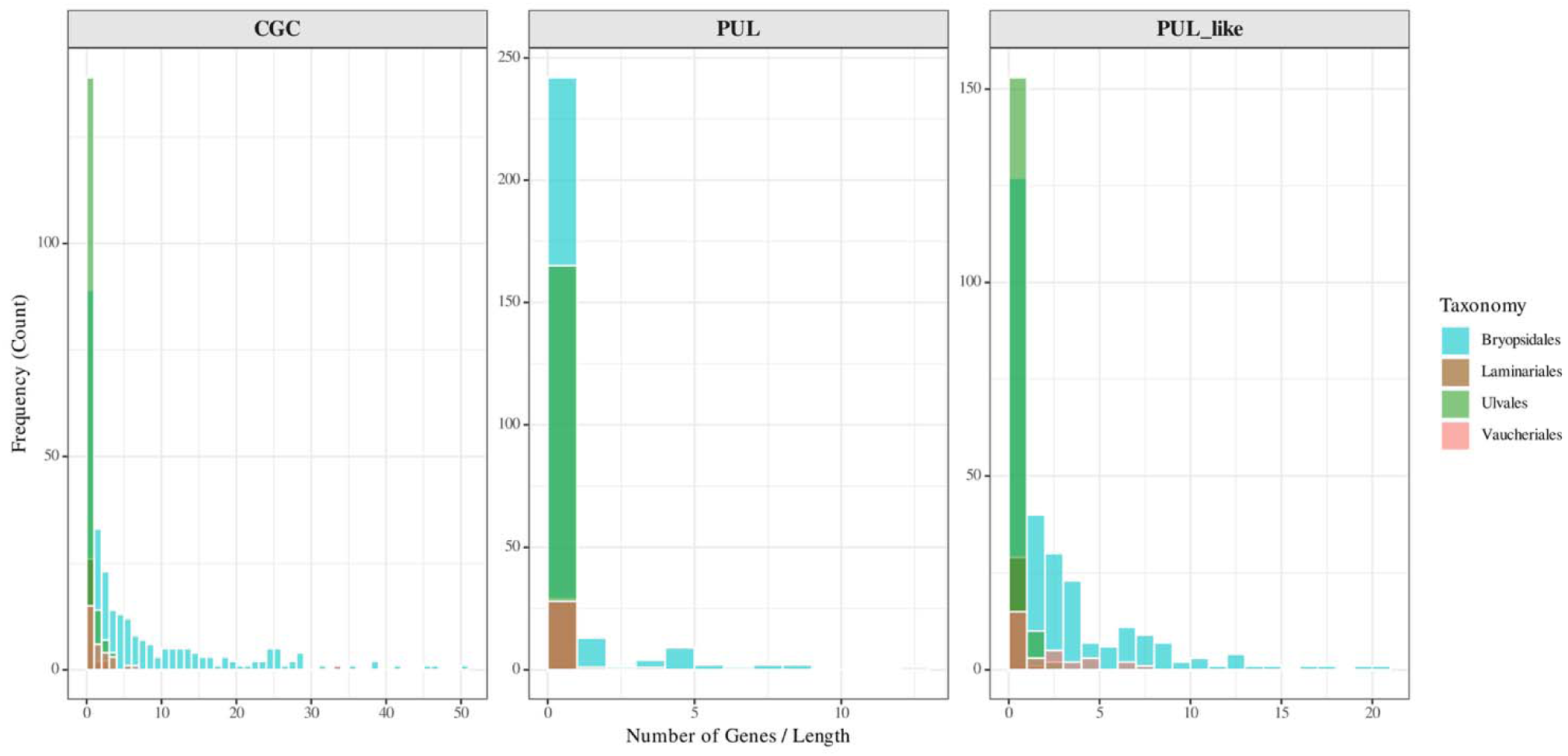
Histogram of the lengths of putative PULs, PUL-like, and CGC found in both MAGs from algae of coenocytic and non-coenocytic orders.

A comprehensive analysis of recovered MAGs from SR and LR of coenocytic algae from the Vaucheriales and Bryopsidales orders showed high dominance at the phylum level of Pseudomonadota. Conversely, in order and family, there is no specific pattern in predicted taxonomy by GTDB-tk. Moreover, this lack of pattern in taxonomic distribution could also be attributable to environmental factors previously observed in the algal sequencing procedure. KEGG pathways in MAGs from Bryopsidales and Vaucheriales showed high completeness in bidirectional polyphosphate metabolism and riboflavin biosynthesis; however, these pathways were significantly more complete in MAGs recovered from non-coenocytic algae (Figure 4A). Conversely, pathways significantly enriched in coenocytic-associated MAGs included protein secretion (Sec-SRP), motility (Flagellum, Chemotaxis), and nutrient provisioning pathways (Thiamin biosynthesis, MEP-DOXP, phosphate transporters), suggesting that the coenocytic morphology could selectively favor bacteria with enhanced host-interaction and biosynthetic capacities. In MAGs from Vaucheriales, their specific pathway prevalence for PHB synthesis that is reported to be a survival mechanism for carbon reserve in low nutrient environments. In addition, the high number of MAGs with 83% completeness of PHB synthesis shows that the *Vaucheria* species could be a potential source for extracting bacteria with this PHB capability, given the nuance of PHB in bioeconomic applications (Verlinden et al., 2007). The NAGGN BGCs cluster is specific and only present in MAGs from algae of the coenocytic order. NAGGN was reported to be associated with a survival mechanism for osmotic pressure. The high completeness of specific KEGG pathways and NAGGN BGCs within MAGs of the coenocytic algae suggests a bacterial community equipped with various survival mechanisms adapted to diverse and challenging environments. Given this observation of the adaptation of survival mechanisms, it can be hypothesized that it may be due to the cosmopolitan distribution of the various coenocytic algae from which MAGs were recovered.

This study represents an initial step toward elucidating the complexity of the microbiome associated with coenocytic algae from the orders Bryopsidales and Vaucheriales, as well as exploring potential associations with host coenocytic morphology.

## Conflict of Interest

*The authors declare that the research was conducted in the absence of any commercial or financial relationships that could be construed as a potential conflict of interest*.

## Author Contributions

GL and AA conceived and designed the study. GL, XR, AS, CJ, CC, YH, and JS performed experiments, analysis, and data collection. GL and XR conducted data analysis and interpretation. GL drafted the manuscript, and AA finalized the manuscript. RRV contributed to bioinformatics analysis. AA supervised the study and secured funding. All authors contributed to the article and approved the submitted version.

## Funding

This study was supported by the National Science Foundation (NSF-HSI STEM; award no. 1928792), Title V DHSI and by CSU Stanislaus Faculty Startup funds awarded to Alok Arun.

## Data Availability Statement

The data set generated in this study was deposited in the NCBI SRA database with the following accession <u>SRX30956066</u>. The sequence of MAGs and functional annotation data is available on figshare: https://doi.org/10.6084/m9.figshare.31953375.

## Notes

https://doi.org/10.6084/m9.figshare.31953375.

### Competing Interest Statement

The authors have declared no competing interest.

