## Supplementary Material for "Genomic Insights into Bacterial Communities of Coenocytic Algae Using Metagenome Assembled Genomes"

### 1 Supplementary Data

**Taxonomy prediction by GTDB-tk**

The taxonomic profiles of the coenocytic orders Bryopsidales and Vaucheriales, determined by GTDB-Tk, display broad similarities at the phylum level but diverge significantly at finer taxonomic resolutions. At the phylum level, both lineages are heavily dominated by Pseudomonadota and Bacteroidota. However, the extent of this dominance varies significantly. In the Vaucheriales microbiome, these two phyla collectively comprise 84.4% of the assemblage, driven by a massive enrichment of Pseudomonadota (64.4%) followed by Bacteroidota (20.0%). The remaining community fraction in Vaucheriales is composed of Bdellovibrionota (8.9%), Actinomycetota (4.4%), Planctomycetota (4.4%), and minor traces of Acidobacteriota (2.2%) and Armatimonadota (2.2%). In contrast, the Bryopsidales microbiome accounts for a lower combined dominance of the top two phyla (72.9%), with Pseudomonadota representing 44.7% and Bacteroidota representing 28.2%. This is distinguished by a significantly higher diversity of associated phyla, including a notable enrichment of Planctomycetota (12.6%), Actinomycetota (3.1%), Myxococcota (2.3%), Cyanobacteriota (1.9%), and Verrucomicrobiota (1.5%), reflecting a more complex holobiont structure.

At the order level, the ecological divergence becomes more pronounced. The MAGs from Vaucheriales were characterized with Sphingomonadales, Burkholderiales, and Rhizobiales each contributing 11.1% (cumulatively 33.3%). This is supported by a secondary tier of surface-associated orders, including Cytophagales and Caulobacterales (both 8.9%), and the predatory order Bacteriovoracales (6.7%). In contrast, the Bryopsidales microbiome is structured by potential symbionts, co-dominated by Rhizobiales and Flavobacteriales (both 11.5%), followed by Rhodobacterales (10.7%) and Pirellulales (7.6%). The high abundance of Flavobacteriales in *Bryopsis* correlates with previous culture-dependent studies identifying Flavobacteriaceae as core facultative endophytes in siphonous algae (Hollants et al., 2011). The Bryopsidales order-level diversity is further sustained by Pseudomonadales (8.0%) and Cytophagales (6.1%), whereas orders such as Sphingomonadales and Burkholderiales are present but markedly less dominant than in the *Vaucheria* dataset.

At the family level, the biological distinction is sharpest. The Vaucheriales retain a consistent taxonomy at the order level, dominated by Sphingomonadaceae (11.1%), Burkholderiaceae (8.9%), and Rhizobiaceae (8.9%). Notably, this microbiome includes Bacteriovoracaceae (6.7%), Caulobacteraceae (6.7%), and Spirosomataceae (4.4%), confirming a community structure driven by predation and rapid nutrient cycling. Conversely, the Bryopsidales display a high abundance of Rhodobacteraceae (10.7%) and Flavobacteriaceae (7.6%), alongside the Planctomycota-derived Pirellulaceae (4.2%) and Cyclobacteriaceae (4.2%). Furthermore, GTDB-Tk classification revealed a significant accumulation of novel taxonomic diversity within the Bryopsidales dataset, identifying 11 putative novel genera. In contrast, the Vaucheriales assemblage yielded fewer novel MAGs, consistent with its composition of well-characterized environmental lineages.

**Metabolic KEGG Pathways Distribution Across Coenocytic and Non-Coenocytic Orders**

Metabolic reconstruction across the coenocytic orders Bryopsidales and Vaucheriales, alongside non-coenocytic algal microbiomes Ulva and Saccharina, revealed universally conserved pathways in all lineages, including amino acid biosynthesis (glycine, lysine, and serine) and glycolysis and the TCA cycle. However, distinct functional architectures differentiated the algae from the coenocytic orders regarding nitrogen metabolism. In the Bryopsidales, *Derbesia* and *Bryopsis* KO-2023 *sp.* encoded complete nitrogen fixation pathways, contrasting with the partial completeness ranging from 0.3 to 0.6 observed in *Penicillus* and *Avrainvillea*, and the genomically reduced profiles of the MAGs from *Ostreobium*. In the Vaucheriales, nitrogen fixation appeared strictly context-dependent, with *Vaucheria bursata* LB2067 exhibiting complete nitrogenase modules co-occurring with Cytochrome bd ubiquinol oxidase and Polyhydroxybutyrate synthesis.

Comparing the community structure of recovered MAGs from coenocytic algae using PCA analysis (Figure S4), there is a relationship consistent with shared KEGG pathways. The majority of coenocytic and non-coenocytic algae MAGs shared a similar functional profile, with the exception of both *Caulerpa lentillifera* and *Caulerpa prolifera*. Conversely, the MAG from *Vaucheria bursata* LB2067 in the upper end of the clustering shows an interesting division from the other coenocytic communities, even its wild counterpart, *Vaucheria bursata*.

**BGCs Distribution Across Coenocytic and Non-Coenocytic Orders**

We evaluated the distribution of Biosynthetic Gene Clusters (BGCs) between the MAGS from the algae of the coenocytic orders Bryopsidales and Vaucheriales and non-coenocytic orders Ulvales and Laminariales to assess any specific traits. Notably, despite lower algae representation in Vaucheriales and Ulvales (2 and 1, respectively), their bacterial communities exhibited a remarkably high density of BGCs relative to the number of algae compared to those in Bryopsidales. When comparing the BGC composition across lineages, the majority of the bacterial community showed high similarity in conserved clusters, specifically Terpenes, Terpene-precursors, and Type I Polyketide Synthases (T1PKS), which were present in both coenocytic and non-coenocytic lineages (Figure 4B).

In contrast, both coenocytic orders were characterized by the specific presence of Butyrolactone and N-acetylglutaminylglutamine amide (NAGNN) gene clusters. Butyrolactone was exclusive to the bacterial community of the Bryopsidales, *Bryopsis* KO-2023 *sp*. NAGNN clusters, while low in total count (n=12), were more broadly distributed among the coenocytic algae. The distribution of NAGNN was highest in *Ostreobium* (33%) and *Derbesia* (32%), followed by the *Bryopsis* genus (16%), and *Vaucheria* and *Penicillus* (8% each).

Finally, both coenocytic algae from the Vaucheriales and Bryopsidales orders, MAGS, shared the usage of the NAGNN cluster. However, the MAGs from the Vaucheriales differentiated themselves through specific, complex gene clusters not found in Bryopsidales. These included phosphonate-like clusters and complex hybrid architectures such as NRP-metallophore-NRPS-T1PKS and T3PKS-terpene hybrids.

### 2 Supplementary Figures and Tables

#### 2.1 Supplementary Figures

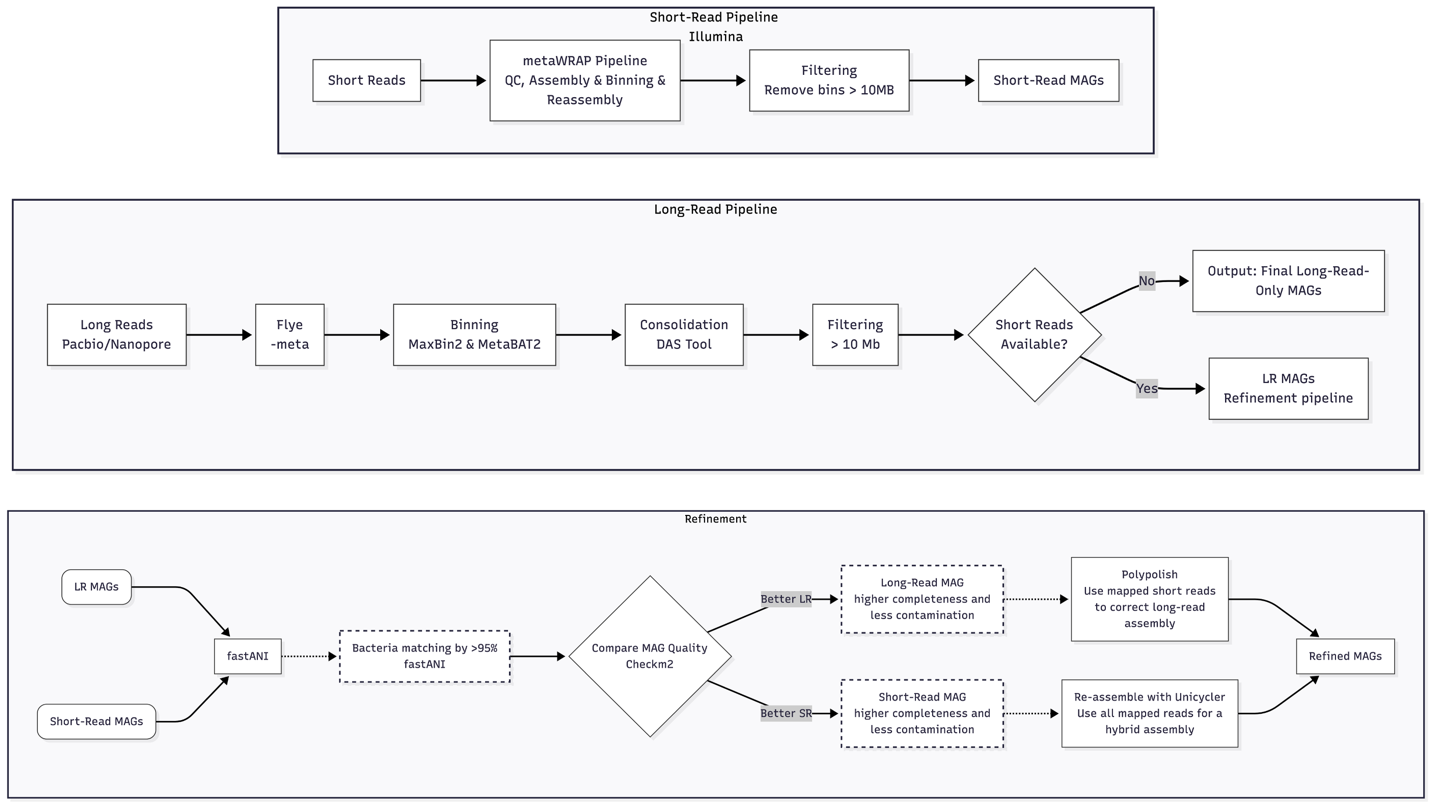

**Figure S1**. Workflow for MAGs recovery from short and long reads, used for the study. This workflow processes long-read and short-read data either separately or in combination to generate initial Metagenome-Assembled Genomes (MAGs). The resulting bins are then compared using quality metrics to select the best version of each genome. Finally, the selected candidates undergo polishing or hybrid re-assembly to produce a refined final dataset.

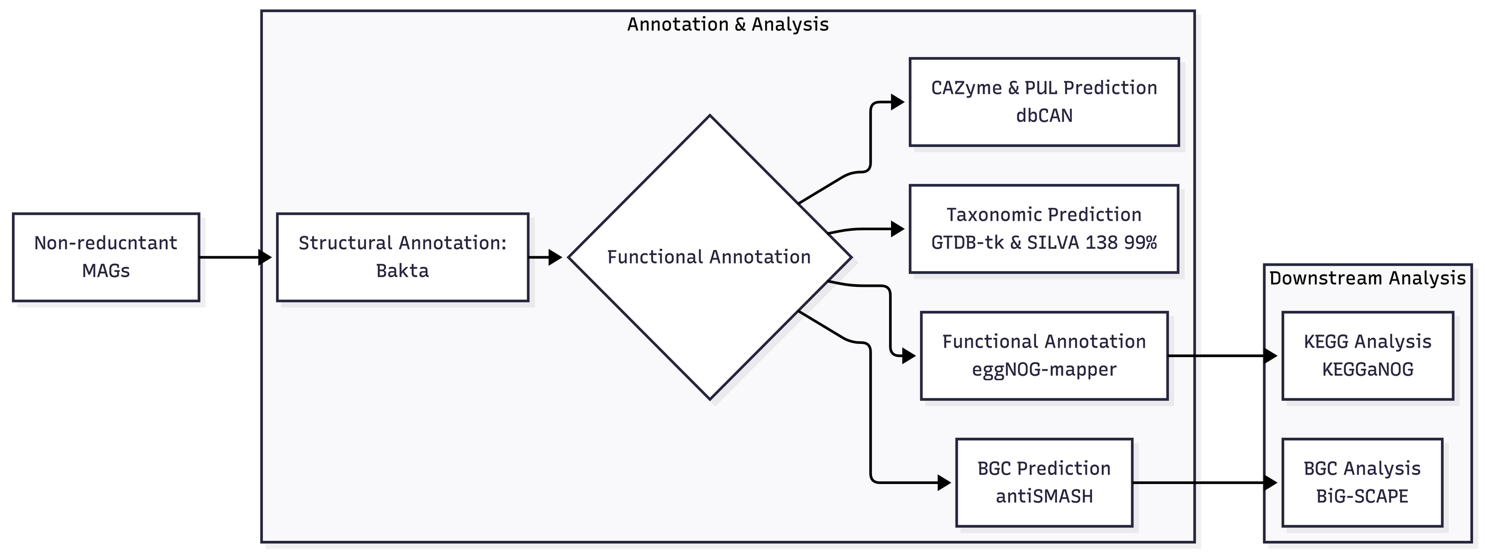

**Figure S2**. Workflow for functional annotation and taxonomical prediction from the recovered MAGs.

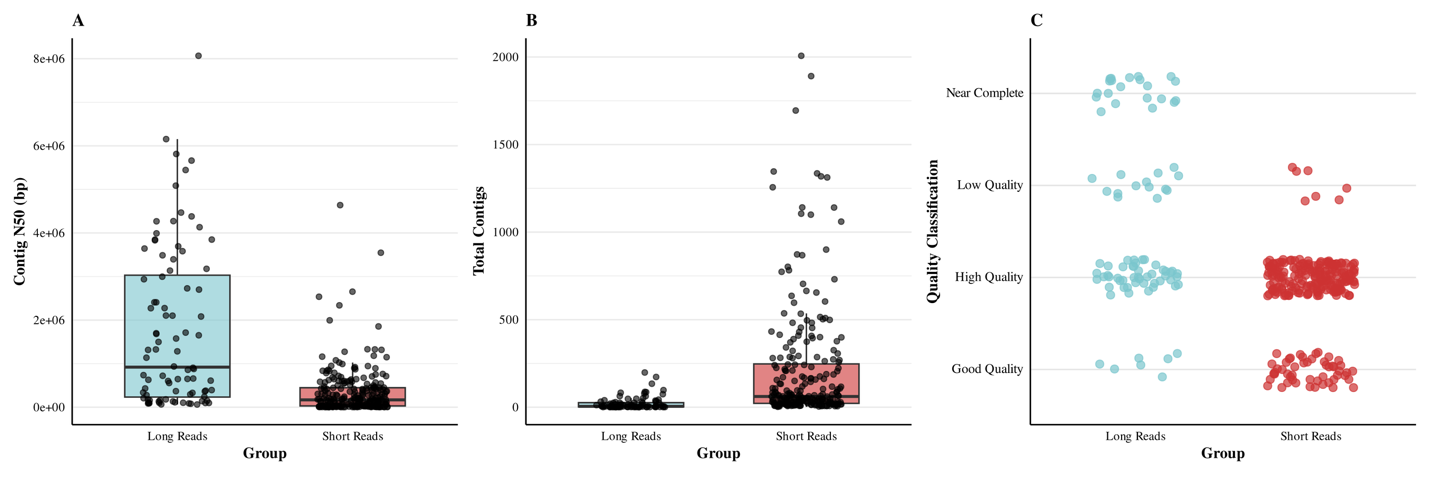

**Figure S3**. Quality metrics of MAGs recovered from long reads (LR) and short reads (SR) from WGS reads of coenocytic algae from the Bryopsidales and Vaucheriales orders. (A) Distribution of N50 of recovered MAGs. (B) Profile of total contigs of MAGs. (C) Categorical distribution of quality across recovered MAGs.

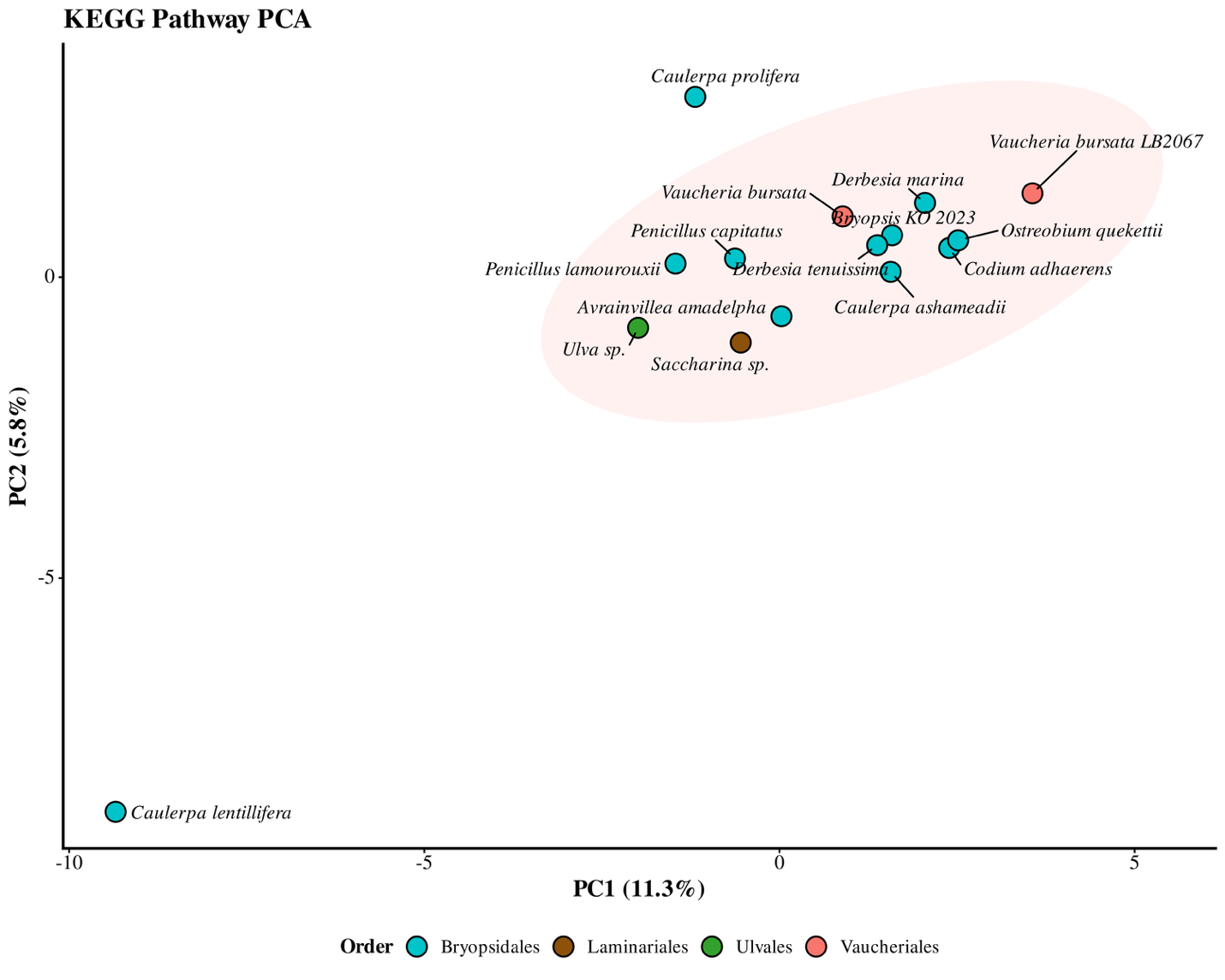

**Figure S4**. Principal component analysis of KEGG pathways profiles from recovered MAGs of coenocytic and non-coenocytic orders. Each point represents a community consensus by calculating by aggregating the KEGG pathways profile of the MAGs from the algae. The shaded ellipse represents a 95% confidence interval. Organisms are colored according to the taxonomic order of the algae.

#### 2.2 Supplementary Tables

**Table S1**. Summary statistics of total MAGs extraction from WGS reads of coenocytic Algae

| **Organism** | **Order Taxonomy** | **Bioproject** | **Total MAGs** | **Near Complete** | **HQ** | **GQ** | **LQ** | **Sequencing Platform** |
| --- | --- | --- | --- | --- | --- | --- | --- | --- |
| *Vaucheria bursata LB2067* | Vaucheriales | [PRJNA1020787](https://www.ncbi.nlm.nih.gov/bioproject/PRJNA1020787) | 11 | 0 | 10 | 1 | 0 | Illumina Pac Bio |
| *Vaucheria bursata* | Vaucheriales | [PRJNA894593](https://www.ncbi.nlm.nih.gov/bioproject/PRJNA894593) | 30 | 0 | 22 | 8 | 0 | Illumina |
| *Ostreobium queketti* | Bryopsidales | [PRJEB39309](https://www.ncbi.nlm.nih.gov/bioproject/PRJEB39309) | 38 | 1 | 34 | 3 | 0 | Illumina  Nanopore |
| *Derbesia tenuissima* | Bryopsidales | [PRJNA924561](https://www.ncbi.nlm.nih.gov/bioproject/PRJNA924561) | 64 | 0 | 43 | 16 | 5 | Illumina |
| *Derbesia marina* | Bryopsidales | [PRJNA924561](https://www.ncbi.nlm.nih.gov/bioproject/PRJNA924561) | 42 | 0 | 38 | 4 | 0 | Illumina |
| *Codium adhaerens* | Bryopsidales | [PRJNA924561](https://www.ncbi.nlm.nih.gov/bioproject/PRJNA924561) | 34 | 0 | 28 | 5 | 1 | Illumina |
| *Caulerpa lentillifera* | Bryopsidales | [PRJDB5734](https://www.ncbi.nlm.nih.gov/bioproject/PRJDB5734) | 4 | 0 | 2 | 2 | 0 | Illumina Pac Bio |
| *Caulerpa ashmeadii* | Bryopsidales | [PRJNA515488](https://www.ncbi.nlm.nih.gov/bioproject/PRJNA515488) | 6 | 0 | 6 | 0 | 0 | *Illumina |
| *Avrainvillea amadelpha* | Bryopsidales | [PRJNA924561](https://www.ncbi.nlm.nih.gov/bioproject/PRJNA924561) | 30 | 0 | 17 | 13 | 0 | Illumina |
| *Penicillus capitatus* | Bryopsidales | [PRJEB87011](https://www.ncbi.nlm.nih.gov/bioproject/1236204) | 61 | 16 | 32 | 5 | 8 | Pac Bio |
| *Penicillus lamourouxili* | Bryopsidales | [PRJEB88235](https://www.ncbi.nlm.nih.gov/bioproject/1248365) | 19 | 4 | 8 | 2 | 5 | Pac Bio |
| *Caulerpa prolifera* | Bryopsidales | [PRJEB94584](https://www.ncbi.nlm.nih.gov/bioproject/PRJEB94584) | 7 | 1 | 5 | 0 | 1 | Pac Bio |

* Illumina data was used instead of long reads from nanopore because of the low quality. High quality (HQ), good quality (GQ), low quality (LQ), and near-complete.

**Table S2**. Quality by CheckM2 of MAGs recovered from Coenocytic Algae

**Table S3**. Taxonomical prediction by GTDB-tk of MAGs recovered from Coenocytic Algae

**Table S4**. Final Taxonomical prediction by GTDB-tk of MAGs recovered from Coenocytic Algae

**Table S5**. Metabolic KEGG pathways of MAGs recovered from coenocytic and non-coenocytic Algae.

**Table S6**. BGCs’ distribution of MAGs recovered from coenocytic and non-coenocytic Algae.

**Table S7**. PUL’s profile of MAGs recovered from coenocytic and non-coenocytic Algae.

**Table S8**. Statistical Analysis of KEGG Pathways of MAGs recovered from coenocytic and non-coenocytic Algae.
