## Supplementary figures and images for "Genomic Insights into Bacterial Communities of Coenocytic Algae Using Metagenome Assembled Genomes"

### Figure S1

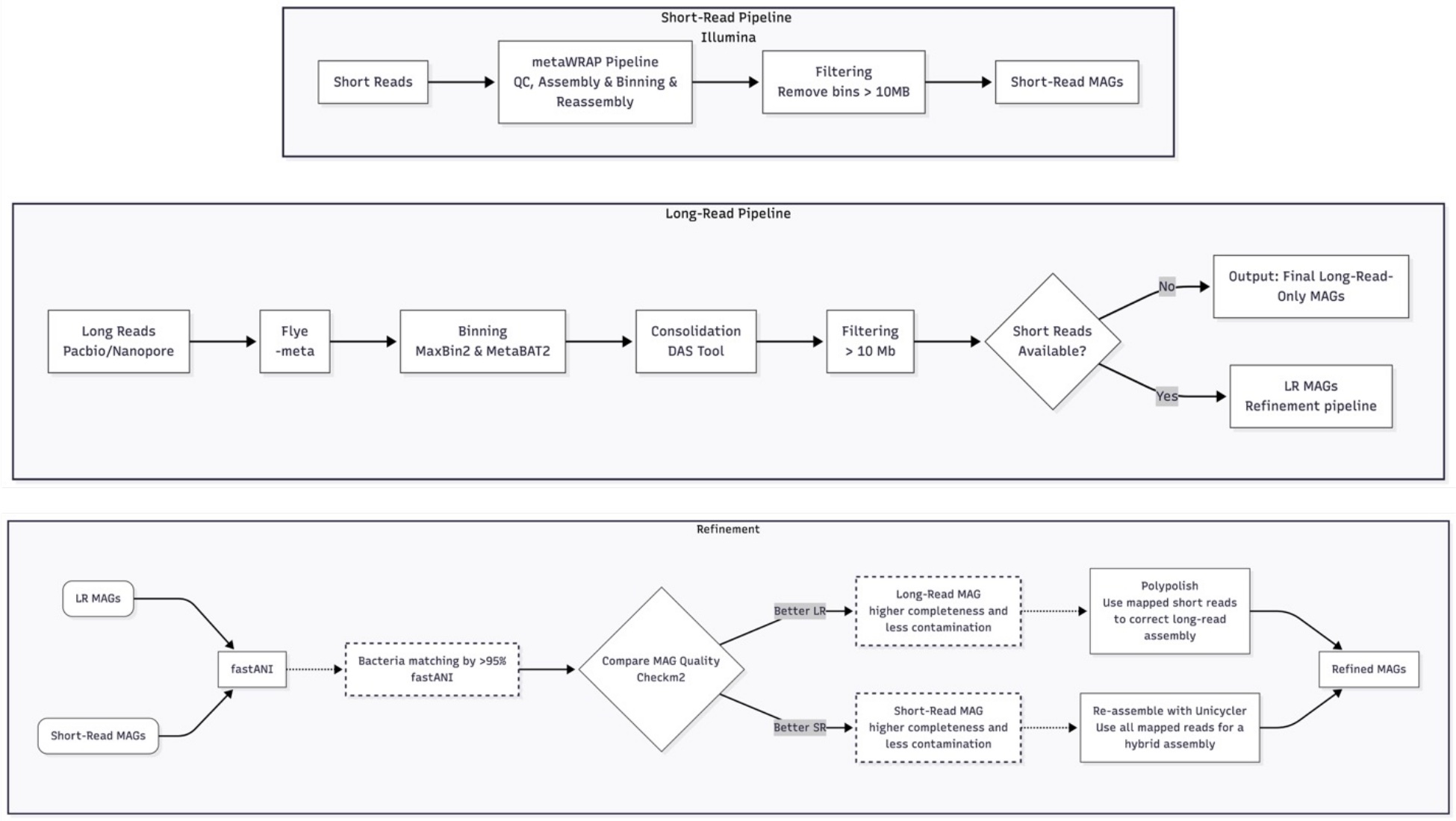

### Figure S2

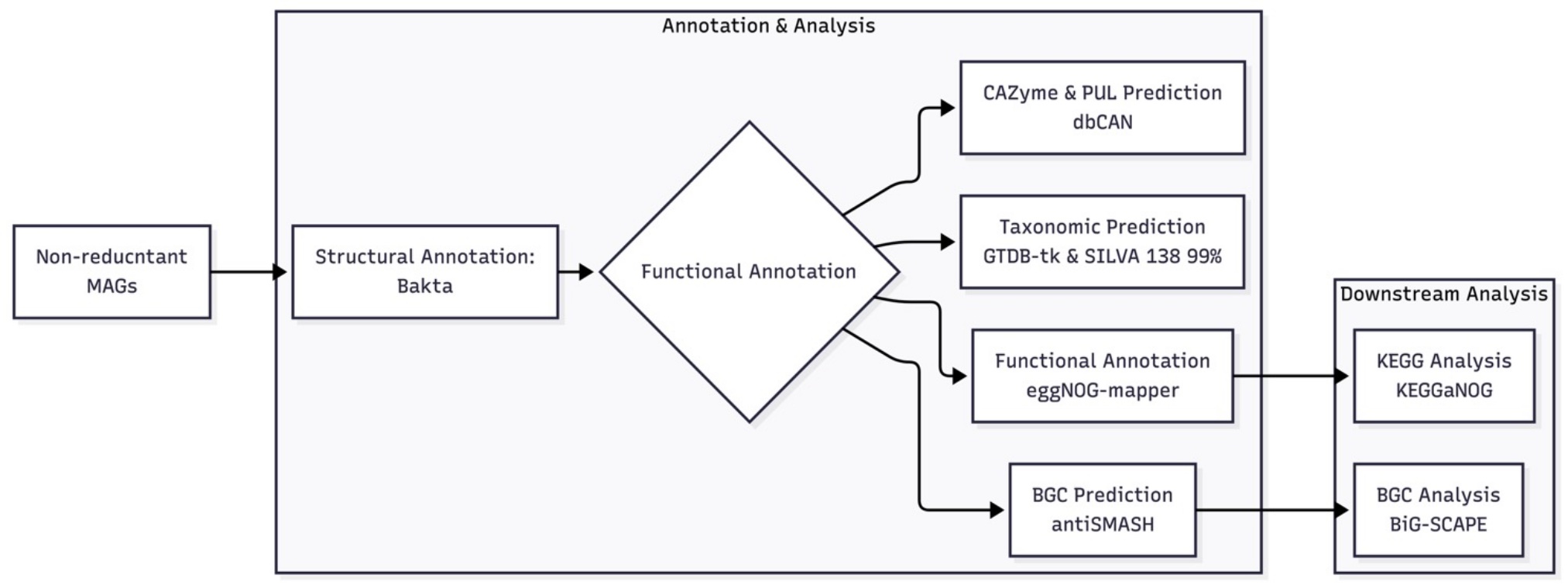

### Figure S3

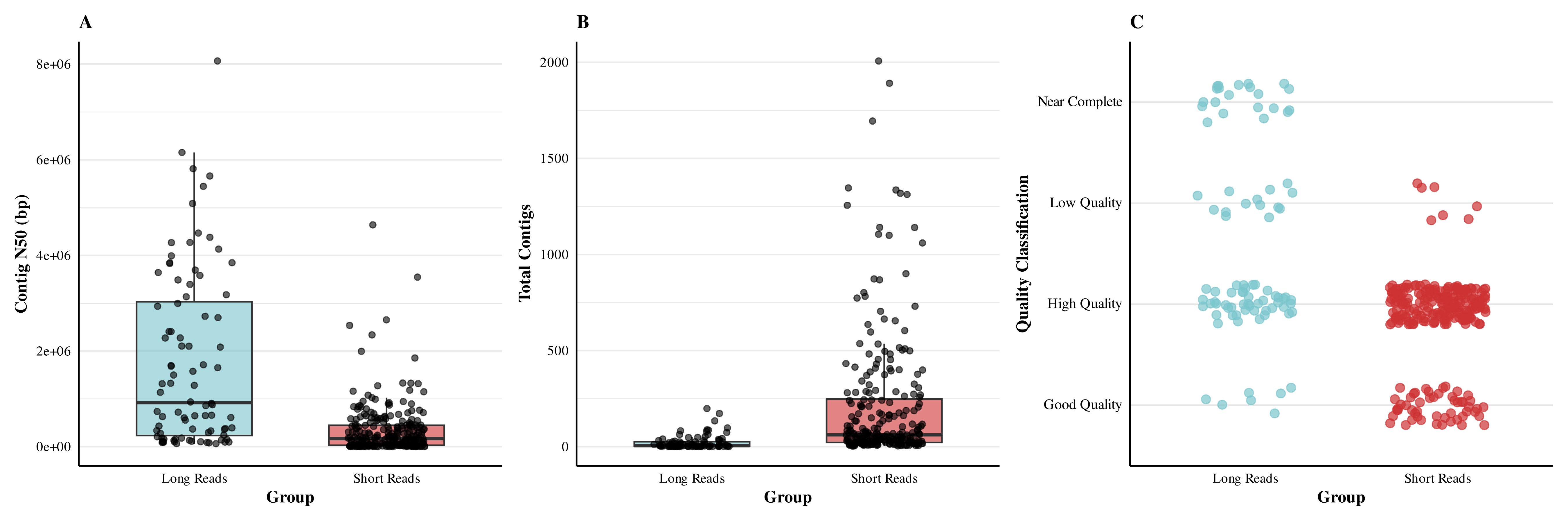

### Figure S4

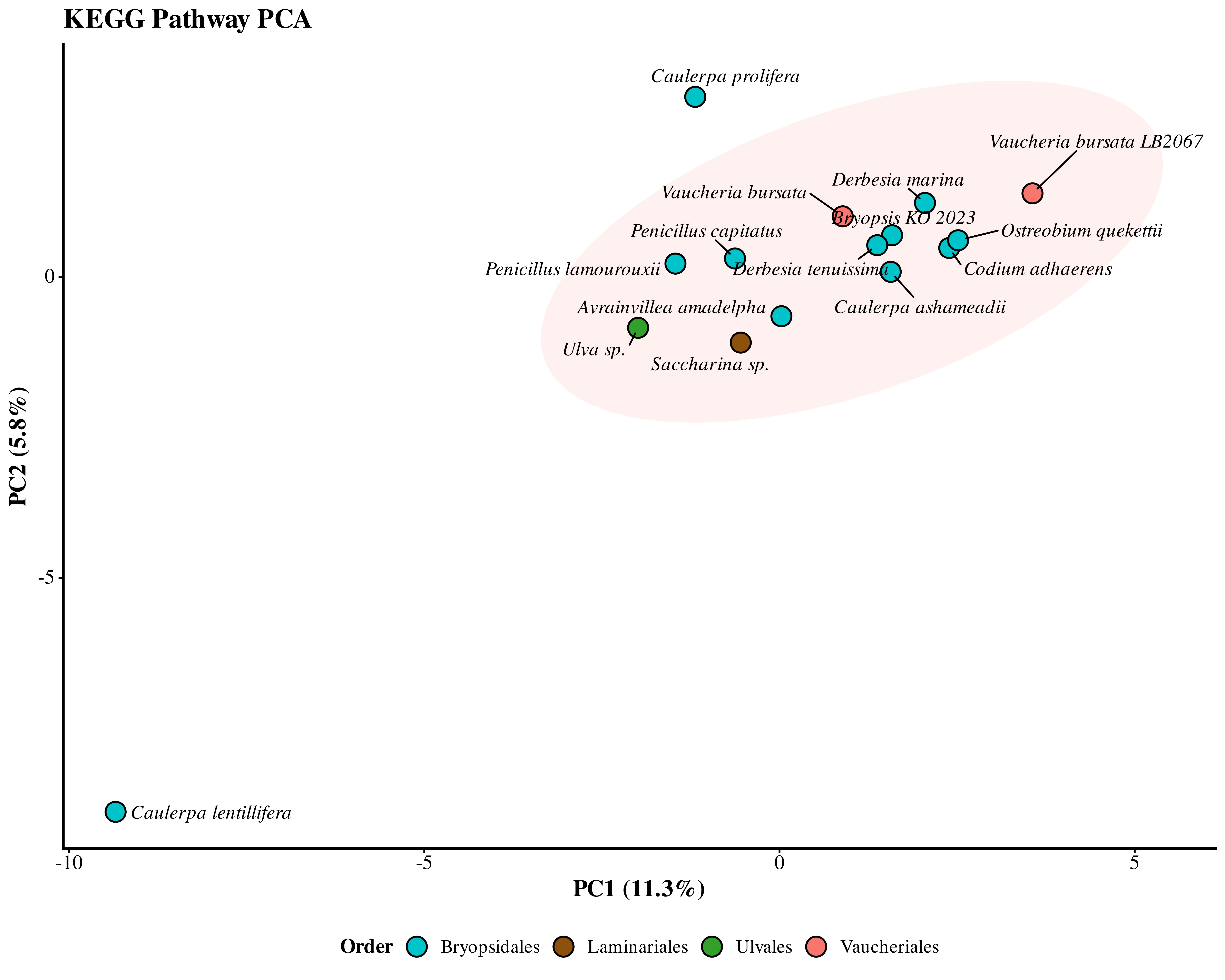
